# Novel anti-repression mechanism of H-NS proteins by a phage “early protein”

**DOI:** 10.1101/2021.03.03.433722

**Authors:** Fredj Ben Bdira, Liang Qin, Alexander N. Volkov, Andrew M. Lippa, Amanda M. Erkelens, Nicholas Bowring, Aimee L. Boyle, Marcellus Ubbink, Simon L. Dove, Remus T. Dame

## Abstract

H-NS family proteins, bacterial xenogeneic silencers, play central roles in genome organization and in the regulation of foreign genes. It is thought that gene repression is directly dependent on the DNA binding modes of H-NS family proteins. These proteins form lateral protofilaments along DNA. Under specific environmental conditions they switch to bridging two DNA duplexes. This switching is a direct effect of environmental conditions on electrostatic interactions between the oppositely charged DNA binding and N-terminal domains of H-NS proteins. The *Pseudomonas* lytic phage LUZ24 encodes the early protein gp4, which modulates the DNA binding and function of the H-NS family protein MvaT of *Pseudomonas aeruginosa*. However, the mechanism by which gp4 modulates MvaT activity remains elusive. In this study, we show that gp4 specifically interferes with the formation and stability of the bridged MvaT-DNA complex. Structural investigations suggest that gp4 acts as an “electrostatic zipper” between the oppositely charged domains of MvaT protomers, and stabilizes a structure resembling their “half-open” conformation, resulting in relief of gene silencing and adverse effects on *P. aeruginosa* growth. The ability to control H-NS conformation and thereby its impact on global gene regulation and growth might open new avenues to fight *Pseudomonas* multidrug resistance.

## Introduction

Bacteriophages - viruses that infect bacteria - are among the most abundant and diverse organisms on earth (1,2) and are found wherever bacteria exist. Bacteria and their associated bacteriophages co-evolve in a continuous battle, each developing defensive and offensive strategies (3-5). To protect themselves against bacteriophages, bacteria have evolved various defense mechanisms, encoded in their genomes, including restriction-modification (6), CRISPR-Cas (7), and xenogeneic silencing systems (8). Bacterial xenogeneic silencers play essential roles in bacterial evolution by recognizing and silencing foreign genes acquired through horizontal gene transfer (9,10), resulting from transformation, conjugation, or transduction. The silencing of these foreign genes, up to the moment that their expression is induced following an environmental cue, can provide bacteria with a competitive advantage under specific conditions without compromising global genome regulation (11).

Four types of bacterial xenogeneic silencers have been identified to date. These proteins belong to the family of H-NS proteins, defined by their functional similarity to the H-NS protein of *E. coli* (8,12,13). Members include H-NS of *Proteobacteria* (14), MvaT and MvaU of *Pseudomonas* species (15), Lsr2 of *Actinomycetes* (16) and Rok of *Bacillus* species (17). The gene silencing mechanism of H-NS family proteins is determined by their ability to bind and spread across genes. Characteristic of H-NS family proteins is the formation of nucleofilaments along the DNA that can switch to mediate DNA-DNA bridges in response to environmental changes (18-23). This interchange between the DNA binding modes of H-NS family proteins is thought to play an important role in the global regulation of gene expression and the dynamic organization of bacterial genomes (24-27).

Bacteriophages have evolved resistance mechanisms by encoding proteins antagonizing H-NS silencing in response to bacterial xenogeneic silencing systems, several of them belong to the phage’s “early proteins”. For instance, the gp5.5 protein of *E. coli* phage T7 can counteract H-NS activity in *E. coli* upon phage infection (28). The gp5.5 binds to the central oligomerization domain of H-NS and disrupts higher-order H-NS-DNA complexes, leading to counter-silencing of genes controlled by H-NS (29,30). Another example is the Arn protein of *E. coli* phage T4, a DNA mimic, which has been proposed to counteract H-NS-mediated repression by targeting its DNA binding domain and thus interfering with DNA binding (31). *E. coli* phage T4 also encodes an “early protein,” termed MotB, that plays a role in improving phage fitness. Its heterologous expression was found to be toxic in *E. coli* B or K12 strains. It was suggested that MotB contains a DNA binding domain with which it can interact tightly, yet non-specifically, with DNA to alter the expression of specific host genes, including the ones repressed by H-NS (32). Bacteriophage Mu employs yet another mechanism. Binding of IHF to its site upstream of the early promoter (Pe) interferes with H-NS-DNA complex formation and counteracts the H-NS-mediated repression of this promoter (33,34). The EBPR podovirus 1 genome encodes a H-NS family protein (gp43), which was proposed to repress expression of genes involved in host defense mechanisms such as CRISPR-associated proteins and a Type III restriction-modification system (35). Regrettably, despite their importance, detailed molecular descriptions of H-NS inhibition by these phage defense mechanisms are generally lacking.

The *Pseudomonas* phage LUZ24 encodes an “early protein,” gp4, that binds the H-NS family protein MvaT (36). MvaT is thought to play an important role in the genome organization of *P. aeruginosa* and in the regulation of gene transcription (15,37). The fold architecture of the MvaT monomer consists of an α-helical N-terminal oligomerization domain (NTD: residues 1-64) tethered to a C-terminal DNA binding domain (DBD) (residues 79-124) by a flexible linker (residues 65-78) (Figure S1b) (18). The structural unit of MvaT protofilaments is a dimer (protomer) formed by “coiled-coil” interactions between the N-terminal α-helices of the MvaT monomers (α1: residues 1-32), dimerization site 1. The MvaT protomers form high-order oligomers through a second dimerization site, site 2 (α2: residues 50-58). Both coiled-coils display a hydrophobic core stabilized by salt bridges (18).

The heterologous expression of gp4 in *Pseudomonas* causes adverse effects on pathogen growth. It was proposed that gp4 inhibits the binding of MvaT to DNA to abolish silencing of the phage LUZ24 genome mediated by MvaT (36). Several possible mechanisms for gp4 modulation of MvaT function can be envisioned based on the MvaT-DNA complex formation pathway determined by the nucleation-propagation-bridging steps (18,19). Therefore, gp4 may alter 1) the DNA binding affinity of MvaT, which is essential for protomer binding; 2) the multimerization properties of MvaT, which determine the cooperativity of nucleoprotein filament formation; or 3) the ability of MvaT to bridge DNA. In this study, we have scrutinized which of these steps in the bridged complex assembly is affected by gp4. Using biophysical and structural biology methods, we defined how gp4 interacts with MvaT and how this translates into altered DNA structuring properties. We propose a molecular mechanism by which gp4 exerts toxicity on *P. aeruginosa in vivo* and may relieve the MvaT-mediated silencing of genes.

## Materials and Methods

### Construction of plasmids

The plasmids encoding MvaT of *P. aeruginosa* (pRD228), its derivatives MvaT F36D/M44D (pRD277), MvaT 1-62-His (pRD360) and MvaT 79-124-His (pRD361)) and gp4 of *Pseudomonas* phage LUZ24 (pRD232) were constructed using pET30b as vector by Gibson assembly (38). The cloned sequences were verified by DNA sequencing.

### DNA substrates

All tethered particle motion and bridging assay experiments were performed using a random, AT-rich, 685 bp (32% GC) DNA substrate used in previous research (18,19). The DNA substrate was generated by PCR and the products were purified using a GenElute PCR Clean-up kit (Sigma-Aldrich). If required, DNA was ^32^P-labeled as described previously (19). For the electrophoretic mobility shift assay an AT-rich 200 bp (31%GC) was generated using the same procedure.

### Protein expression and purification

BL21 (DE3) pLysS cells transformed with pRD228, pRD277, pRD360, pRD361 or pRD232 were grown in 2l of LB supplemented with kanamycin (100 µg/ml) at 37°C to an OD600 of ∼0.6. Protein expression was induced with 0.5 mM isopropyl β-D-1-thiogalactopyranoside (IPTG) overnight at 18°C. Cells were centrifuged at 7000 × g for 15 min at 4°C. MvaT and MvaT F36D/M44D were purified as described previously (18). MvaT 1-62-His and MvaT 79-124-His were purified with modifications. The harvested cells were lysed by sonication in a lysis buffer containing 20 mM Tris-HCl (pH 8.0), 1 M NaCl. The lysate was cleared by centrifugation at 37 000 × g for 30 min at 4°C. Next, the supernatant was loaded on a HisTrap HP 5ml column (GE healthcare Life sciences) and the protein was eluted by applying an imidazole gradient from 0 to 1 M. The eluted fractions were checked by SDS-PAGE and the fractions that contain the target protein were pooled, concentrated and buffer exchanged using a PD10 column to 20 mM tris-HCl pH 7, 100 mM KCl. Next, the MvaT 1-62-His and MvaT 79-124-His were loaded on a HiTrap Q HP 5ml and HiTrap SP HP 5ml column (GE healthcare Life sciences) respectively and eluted by applying a NaCl gradient (from 0.1 to 1 M). The eluted fractions were also checked by SDS-PAGE and the pooled fractions were concentrated to a 500 μl volume with an Amicon 10 kDa cut-off filter. The concentrated protein fractions were loaded on a GE Superdex 75 10/300 GL column and eluted with 20 mM Tris-HCl pH 8, 300 mM KCl, 10% Glycerol. For the purification of gp4, the harvested cells were lysed by sonication in a lysis buffer containing 20 mM Tris-HCl (pH 7.0), 100 mM NaCl. Next, the supernatant was loaded on a HiTrap SP HP 5ml column (GE healthcare Life sciences) and the protein was eluted by applying a NaCl gradient from 0.1 to 1 M. The eluted fractions were checked by SDS-PAGE and the fractions that contain the target protein were pooled, concentrated and loaded into a GE Superdex 75 10/300 GL column and eluted with 20 mM Tris-HCl pH 8, 300 mM KCl, 10% Glycerol. The purity of the protein was checked by SDS-PAGE and the concentration was determined using a Pierce BCA protein assay kit (Thermo Scientific).

### His-tag pull-down assay

120 µl of the High-Density Nickel Agarose (Jena Bioscience) suspension was pipetted into an Eppendorf tube. The beads were prepared by changing the solution to binding buffer (100 mM NaCl, 10 mM tris-HCl pH 7.2, 50 mM imidazole, 20 mM MgCl_2_). The nickel beads were then centrifuged at 7000 rpm for 2 minutes and the supernatant was removed. 250 µl of the desired combination of MvaT 79-124-His (40 µM), His-MvaT 1-62 (40 µM) and/or gp4 (40 µM), in binding buffer, was added to the prepared nickel beads. This was incubated for 30 minutes while being shaken at 1000 rpm. After incubation the samples were centrifuged at 3000 rpm for 2 minutes. The supernatant was removed, and the pellet was carefully resuspended in 250 µl of binding buffer. This was referred to as a washing step and was repeated a total of six times. Following the final washing step, 250 µl of elution buffer (100 mM NaCl, 10 mM tris HCl pH 7.2, 1.5 M imidazole, 20 mM MgCl_2_) was added to the beads instead of binding buffer to elute any protein bound to the bead. The samples were centrifuged at 3000 rpm for 2 minutes and the supernatant was collected. 10 µl of the supernatant was run on a tricine gel.

### Tethered Particle Motion

Tethered Particle Motion experiments were performed as described previously (18,39) and the flow cells were prepared with minor modifications. In short, first, the flow cell was washed with 100 ml of wash buffer (10 mM Tris-HCl pH 8, 150 mM NaCl, 1 mM EDTA, 1 mM DTT, 3% Glycerol, 100 µg/ml BSA (ac)) and experimental buffer (10 mM Tris pH 8, 50 mM KCl, 10 mM EDTA, 5% glycerol). The flow cell was sealed after adding protein (MvaT alone or MvaT-gp4 mixure) in experimental buffer, followed by 10 minutes of incubation. The measurements were initiated 15 min after introducing protein. For each flow cell more than 200 beads were measured at a temperature of 25°C. Two separate flow cells were measured for each concentration.

### Electrophoretic mobility shift assay

The assay was performed on an AT-rich 200 bp DNA substrate (31% GC). DNA (25 ng) was incubated with different concentrations of the desired combination of MvaT, MvaT F36D/M44D and gp4 in the binding buffer (10 mM Tris-HCl, 60 mM KCl, pH 8.0). The mixture was loaded on a 1% agarose gel (containing 1:104 dilution of Gel red DNA stain) with loading buffer (10 mM Tris-HCl, pH 7.6, 0.03% bromophenol blue, 0.03 % xylene cyanol FF, 60% glycerol and 60 mM EDTA). The samples were run at 4°C in TAE standard buffer. The gels were visualized with a Bio-rad Gel Doc XR+ system.

### Isothermal Titration Calorimetry

Isothermal Titration Calorimetry experiments were performed as described previously (18) with modifications. Briefly, ITC experiments were performed using a MicroCal VP-ITC system at 20 °C. The protein samples were dialyzed to a buffer containing 20 mM Bis-Tris, pH 6, 50 mM KCl, 16 mM MgCl_2_. Typically, 20 µM of MvaT F36D/M44D was placed in the cell (1.4 ml) and titrated with 300 µM (500 µl) of the gp4, injected in 2 µl aliquots. The delay time between the injections was 60 s with a stirring speed of 307 rpm. The corresponding “protein to buffer” controls were performed for background correction. The ITC titration data were analyzed using Origin 7.0 (OriginLab) provided with the instrument. One set of sites model (1:1 interaction) was used to fit the data where N (number of binding sites), K (association constant) and ΔH (delta enthalpy) were set free during fitting.

### Bridging assay

The bridging assay was performed as described previously (18,39) with modifications. To start the bridging reaction, MvaT was added, followed by 2 µl of gp4 to different final concentrations. The samples were incubated while shaking at 1000 rpm at 25°C. The reactions were stopped and washed at certain time with buffer (10 mM Tris-HCl, pH 8, 65 mM KCl, 5% glycerol, 1 mg/ml BSA (ac), 1 mM spermidine, 20 mM CaCl_2_, 0.02% Tween 20), followed by resuspension in 12 µl denaturing buffer (10 mM tris pH 8, 200 mM NaCl, 1 mM EDTA, 0.2% SDS). Scintillation was used to determine the final radioactivity of each sample. 2 µl of radioactive DNA was used as a reference to calculate the bridging efficiency (DNA recovery %).

### Heterologous expression of LUZ24 gp4 in Pseudomonas aeruginosa

Wild-type *Pseudomonas aeurigonsa* PAO1 was obtained from Arne Rietsch, Case Western Reserve University. The strain harboring a deletions of *mvaU* was previously described (40). The *mvaT* deletion strain was constructed using the pEXG2-based plasmid pEXG2M4315 by allelic exchange (41). The deletion retains the start codon for *mvaT* and replaces the remaining coding sequence with DNA encoding 3 alanines followed by a stop codon. Rhamnose-inducible expression of gp4 was achieved by integration of the pJM220-based plasmid pAL63 in single copy at the Tn7 attachment site of PAO1, the Δ*mvaU* strain, or the Δ*mvaT* strain, as previously described (42) followed by removal of the *aac1* casette using pFLP2 (43).pAL63 includes the coding sequence of gp4 inserted between the SpeI and HindIII sites of pJM220 via isothermal assembly. Freshly streaked colonies of each strain grown on lysogeny broth (LB) agar plates were resuspended in PBS, OD_600_ was determined using a NanoDrop ND1000 spectrophotometer, normalized to OD_600_=0.01 (designated dilution 10^−1^) and serially diluted 10-fold in PBS. 10 µL of culture for each strain and dilution were plated to LB or LB supplemented with L-Rhamnose (Sigma-Aldrich) to a final concentration of 1%. Plates were incubated at 27°C for 18 hours and photographed under white light illumination using a Nikon D3400 camera.

### NMR titration

The NMR titration of the MvaT_2_ with gp4 was performed on a 150 µM (of MvaT monomer) ^15^N isotopically labelled protein sample, in 20 mM Bis-Tris Buffer, 50 mM KCl, 16 mM MgCl_2_, pH 6 and 6% D_2_O. A series of ^1^H-^15^N HSQC spectra were acquired by gradually increasing the MvaT:gp4 molar ratio up to 1:1.2. The experiments were recorded at 20 °C on a Bruker Avance III (HD) 600 MHz spectrometer, equipped with TCI cryoprobe, processed by TopSpin 3.5 (Bruker Biospin) and analyzed by Sparky software (44). The same experimental procedure was applied for the titration MvaT-gp4 (1:1.2 molar ratio) with 20 pb of DNA. The DNA: MvaT-gp4 molar ratio was increased from 0.2 to 1.2. The changes in peak positions and intensities were analyzed and the average chemical shift perturbations (CSP) were calculated using equation (1):

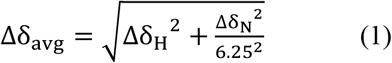

### NMR solution structure of LUZ24 gp4

All NMR experiments were performed at 293 K on a Bruker Avance III HD 800 MHz spectrometer equipped with a TCI cryoprobe. The sample contained 1 mM U-[^13^C, ^15^N] labeled gp4 in 20 mM Bis-TRIS pH 6.0, 150 mM KCl, 1 mM EDTA and 6 % D_2_O for the lock. All NMR data were processed in TopSpin 3.6 (Bruker) or NMRPipe (45) and analyzed in CCPN (46). Nearly complete, unambiguous ^1^H, ^13^C and ^15^N resonance assignments of the protein nuclei were obtained from a suite of standard multidimensional NMR experiments: 2D [^1^H,^15^N]-HSQC, [^1^H,^13^C]-HSQC, and constant-time [^1^H,^13^C]-HSQC for the aromatic region; triple-resonance HNCACB, HN(CO)CACB, HNCO, HN(CA)CO, HBHA(CO)NH, (H)CCH-TOCSY, and H(C)CH-TOCSY experiments; 2D (HB)CB(CGCD)HD and (HB)CB(CGCDCE)HE spectra for the aromatic resonances; and 3D ^15^N-edited NOESY-HSQC and ^13^C-edited NOESY-HSQC for aliphatics and aromatics. The resonance assignments were deposited in the BMRB data bank under the accession number 28112.

The 3D ^15^N-edited NOESY-HSQC and ^13^C-edited NOESY-HSQC spectra for aliphatics and aromatics, all acquired with the mixing time of 120 ms, were subsequently used for the protein structure calculation. The NOE cross-peaks, determined with CCPN Analysis (46), were combined with the dihedral angle restraints, obtained with DANGLE (47), and used as an input for the automated NOE assignment and structure calculations in CYANA v.3 (48), followed by the explicit solvent and torsion angle refinement in CNS (49) and Xplor-NIH (50), respectively. The 10 lowest-energy structures were retained and deposited in the PDB bank under the accession code 6YSE. The NMR structure calculation and refinement statistics are presented in Table S1.

### Modelling of MvaT_2_-gp4 complex

Prediction of the MvaT_2_-gp4 dimer complex was performed by HADDOCK2.2 server (51) using default setting parameters. The data from the NMR titration of the MvaT_2_ with gp4 was used to restrain the docking. The homology model of MvaT dimer (18) was used. Only the amino acids of site 1 of the MvaT dimer were classified as *active* in the docking settings. As most of gp4 amino acid residues appear to be are involved in the complex formation, the full protein amino acid sequence was classified as *active*. Amino acid residues indirectly involved in complex formation (*passive*) were automatically identified by the server. HADDOCK clustered 138 structures in 16 clusters, which represents 69.0 % of the water-refined models HADDOCK generated. The statistics of the top cluster are shown in Table S2. The top cluster is the most reliable according to HADDOCK. The reported scores and energies are averages calculated over the top four members of a cluster.

### Synthesis of gp4 K42E/K45E

The gp4 K42E/K45E protein was synthesised via solid-phase peptide synthesis (SPPS) using a microwave-assisted Liberty Blue peptide synthesiser. A Wang resin, preloaded with the C-terminal lysine residue was utilized as the solid support. Standard Fmoc-chemistry protocols were employed, with Fmoc-deprotection achieved using 20% piperidine in DMF, and coupling reactions facilitated by DIC/Oxyma as the activator/activator base. After synthesis, the protein was cleaved manually from the solid support using a TFA:TIPS:H2O, (95:2.5:2.5) mixture. The protein was subsequently precipitated into ice-cold diethyl ether, collected by centrifugation and freeze-dried. Purification was performed by high-pressure liquid chromatography (HPLC) using a Shimadzu system comprising two KC-20AR pumps and an SPD-20A UV-Vis detector, fitted with a Kinetix Evo C18 column. The protein was eluted from the column using a linear gradient of 20-80% buffer B (A = H2O with 0.1% TFA, B = MeCN with 0.1% TFA) over 20 minutes. The purified protein was freeze-dried and TFA residues were removed by dialysis before the protein was utilized. TFA removal was monitored by 19F NMR.

## Results

### LUZ24 gp4 is selectively targeting MvaT *in-vivo* to modulate its DNA binding modes

Previous studies have suggested that the LUZ24 gp4 protein negatively affects *Pseudomonas* growth by targeting its H-NS family protein, MvaT (36). To validate these findings, we first heterologously expressed gp4 in a wild type strain of *Pseudomonas aeruginosa* (see M&M). Induction of gp4 expression by 1% rhamnose caused severe inhibition of bacterial growth (Figure S1a). However, the expression of gp4 in cells lacking MvaT abrogated that toxicity. MvaU is an MvaT paralog that binds to many of the same genomic regions of *P. aeruginosa* and coordinately regulates the expression of ≈350 target genes (15,37). However, gp4 remained toxic in cells lacking MvaU. These data show that gp4 is toxic to cells of *P*. *aeruginosa* in a manner that is dependent upon MvaT but not MvaU.

The mechanism underlying gp4 activity was earlier proposed to be inhibition of MvaT binding to DNA, based on the results from an electrophoretic mobility shift assay (EMSA) (36). To verify the reported results, we also performed an EMSA using a 200 bp DNA (30% GC) substrate. In our hands, the addition of gp4 from 1 to 4 µM to the DNA in the presence of 2 µM MvaT, does not inhibit the formation of MvaT-DNA complexes (Figure 1a). This discrepancy in results might stem from the use of gp4-streptavidin and 6×His-MvaT tagged proteins in the previous studies, whereas we used untagged proteins.

**Figure 1:**
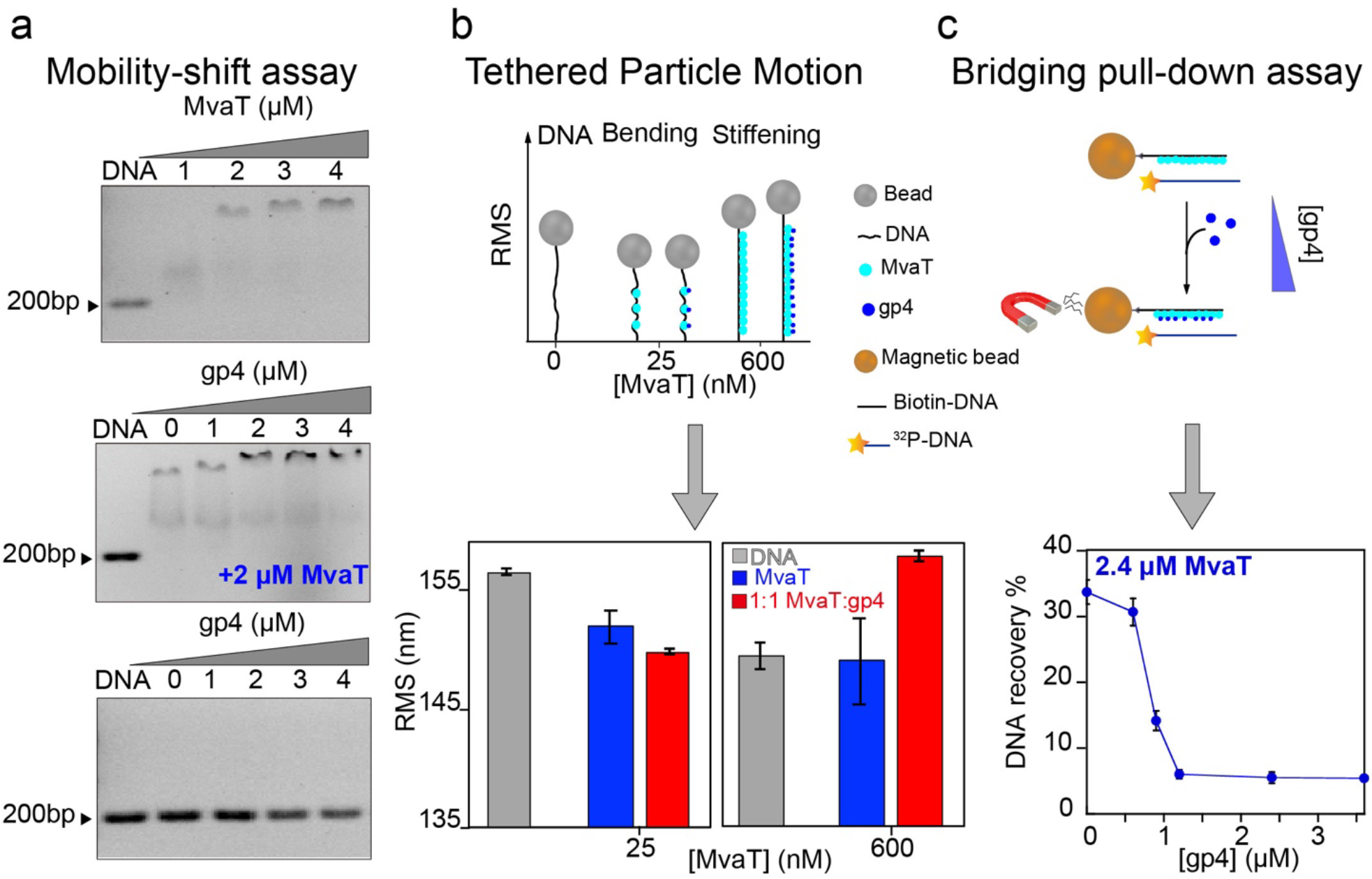
Effect of LUZ24 gp4 protein on the DNA binding modes of MvaT. (a) Electrophoretic mobility-shift assay (EMSA) of 200 bp DNA substrate (32% GC) by MvaT in the absence (upper panel) and presence (middle panel) of different concentrations of gp4. The lower panel represents a control EMSA in the presence of gp4 only. Note the decrease in the DNA band intensity at high concentration of gp4, which might be due to the formation of a non-specific and weak complex with the DNA. (b) & (c) Effects of gp4 on MvaT DNA stiffening and bridging activities, respectively. In the upper panels a schematic representation of the TPM and bridging assays is depicted (see M&M). The lower panels represent the experimental data. The error bars are for the standard deviation from duplicate experiments.

To further validate our observations, we also examined the effect of gp4 on the DNA binding of MvaT F36D/M44D (MvaT_2_), a mutant that exists exclusively as dimer and does not oligomerize further (18). Similar to MvaT WT, the DNA binding of MvaT_2_ was not inhibited by adding increasing amounts of gp4, as concluded from the EMSA results (Figure S1c). Previously, the ^1^H-^15^N HSQC NMR spectrum of MvaT_2_ was assigned and used to study the protein interactions with a 20 bp DNA substrate at low and high ionic strength (18). Here we performed a NMR titration of a ^15^N-MvaT_2_ sample in the presence of gp4 (MvaT:gp4 1:1.2 ratio, see below) with a 20 bp DNA substrate. The titration data show progressive chemical shift perturbations (CSP) of the DNA binding domain (DBD) resonances (Figure S1c), similar to the previous results obtained from the titration of MvaT_2_ with DNA in the absence of gp4 (18). Based on these results, we conclude that gp4 does not inhibit the binding of MvaT to the DNA substrate.

Oligomerization of MvaT is required for the formation of nucleoprotein filaments along DNA and is essential for the function of MvaT in gene silencing (20,52). To investigate the effect of gp4 on the formation of MvaT nucleofilaments, we used Tethered Particle Motion (TPM) (18,19). In the absence of gp4, DNA compaction (indicated by a reduction in the root mean square displacement (RMS)) was observed at a low concentration of MvaT (25 nM). This effect was attributed to DNA bending by individually bound MvaT dimers (nucleation step) (18). Upon increasing the amount of MvaT, stiffening of the DNA substrate is observed, and saturation is reached at 1200 nM MvaT (Figure S1c).

gp4 alone does not alter the RMS, attributed to lack of DNA binding (Figure S1e), in line with the EMSA results (Figure 1a). To test whether gp4 affects the formation of the MvaT-DNA filament, TPM was performed with fixed MvaT concentrations of 25 nM and 600 nM, in the presence of equivalent concentrations of gp4. At 25 nM MvaT, addition of gp4 slightly reduces the RMS of the MvaT-DNA complex (Figure 1b). This indicates that gp4 exhibits minor effect on the bending capacity of the MvaT protomers. At 600 nM MvaT, the MvaT-DNA nucleofilament is not abolished by gp4 (Figure 1b). On the contrary, the increase in RMS indicates that addition of gp4 leads to a more extended MvaT-DNA nucleofilament. Based on the TPM data, we conclude that gp4 does not interfere with the oligomerization of MvaT along DNA and thus does not inhibit its stiffening activity.

In conjunction with DNA stiffening, MvaT can bridge two DNA duplexes under specific buffer conditions (18,19,53). To determine whether gp4 affects the ability of MvaT to bridge DNA, we used a protein-DNA pull-down bridging assay (19,39). MvaT was found to bridge DNA in the presence of divalent or monovalent ions, in a concentration-dependent manner (18). Here we performed the experiments in the presence of 16 mM MgCl_2_, a condition at which MvaT nucleofilaments exhibit optimal DNA bridging activity. A fixed amount of MvaT monomer (2.4 µM) is used, yielding a 35% recovery of DNA in the absence of gp4. The DNA recovery decreased upon addition of gp4. Maximum inhibition of DNA recovery was achieved at a gp4 concentration of 1.2 µM, where the MvaT:gp4 molar ratio is 2:1 (Figure 1c). These results demonstrate that gp4 selectively inhibits the formation of MvaT-DNA bridges, and does not perturb MvaT-DNA filament formation.

Next, to examine whether gp4 affects preassembled MvaT-DNA bridges, we performed a time-dependent DNA bridging assay at a 1:1 molar ratio of MvaT:gp4 (Figure S1f). As a control, we measured the time dependence of DNA recovery by MvaT in the absence of gp4. Under this condition, DNA recovery increases over time and reaches saturation at t = ∼ 600 s. When gp4 was introduced at the beginning of the assay (t = 0 s), a minimal increase in DNA recovery was observed (∼ 5%). Upon addition of gp4 at t = 180 s (pre-saturation level), and 1200 s (at saturation level), a reduction in DNA recovery by about 10% is observed. These data demonstrate that gp4 can perturb, yet not completely disrupt, preassembled MvaT-DNA bridges.

Collectively, we have demonstrated that gp4 specifically targets MvaT in *Pseudomonas* to selectively inhibit its DNA bridging activity. This process is neither caused by inhibition of the DNA binding capacity of MvaT nor by the obliteration of its oligomerization along DNA.

### Structural characterization of LUZ24 gp4

The LUZ24 gp4 is a 46 amino acid peptide with no reported tertiary structure. The secondary structure of the gp4 polypeptide chain is predicted to consist of two α-helices: a long N-terminal (α1: residues 4-26) and a short C-terminal (α2: residues 31-45) helix, connected by a loop of four amino acids (Figure 2a). In agreement with these predictions, gp4 exhibits a far-UV CD spectrum with two minima at approximately 208 and 222 nm, typical of α-helical secondary structures (Figure 2b). The ratio between the intensities of the 222 and 208 nm bands (I_222_/I_208_) is equal to 1.13, suggesting that the peptide is assuming a coiled-coil structure (54). In contrast to the narrow dispersion of resonances, often observed in the ^1^H-^15^N HSQC spectra of α-helices (centred at ∼ 8 ppm), gp4 has a well-dispersed HSQC spectrum (Figure 2c). This characteristic of the NMR spectrum reflects a non-homogeneous chemical environment of the backbone amide nuclei, possibly due to the coiled-coil fold of the peptide.

**Figure 2:**
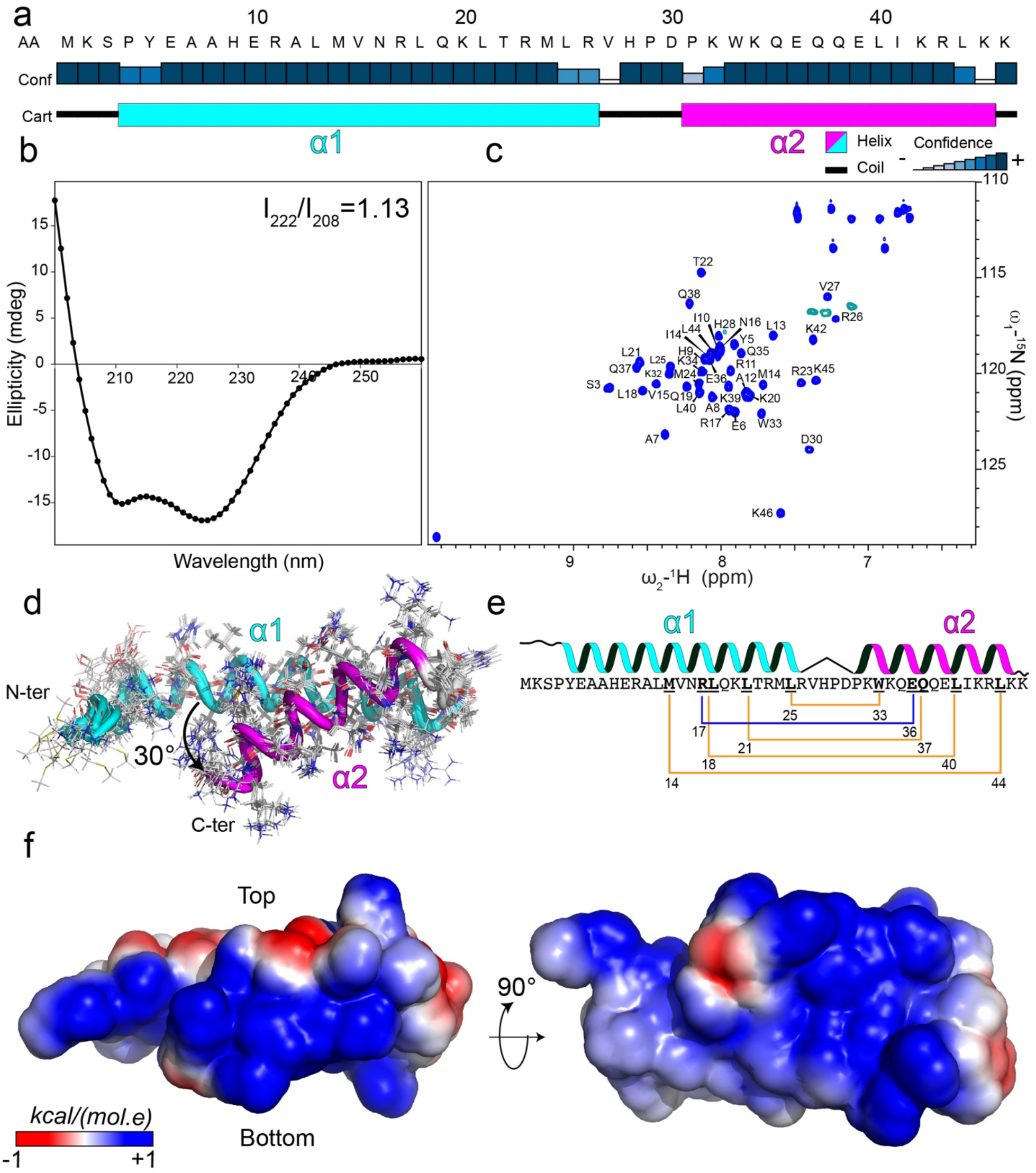
Structural characterization of LUZ24 gp4. (a) Analysis of gp4 secondary structure by JPred 4 server (55). The blue rectangles indicate the confidence of the prediction. The predicted N-terminal helix (α1) is shown in a cyan rectangle and the C-terminal helix (α2) is in magenta rectangle. (b) Far UV-CD spectrum of gp4 recorded at 5 μM protein concentration in 10 mM phosphate buffer, pH 7. The ratio between the 222 nm and 208 nm bands is indicated. (c) Assignment of the ^1^H-^15^N HSQC spectrum of gp4. (d) Bundle of the best 10 structures of gp4 is shown in cartoon and the amino acids side chains in lines. The α−helices are colored as in (a). The angle between gp4 helices is indicated. (e) Schematic representation of the intramolecular interaction within the gp4 coiled-coil fold. The hydrophobic interactions between the side chains of the amino acid residues (bold underlined) are shown with a yellow line and the salt bridge in blue. (f) The electrostatic surface of gp4 is shown, generated by CHARMM-GUI (56).

We determined the solution structure of gp4 using NMR spectroscopy, and the bundle of the ten best structures is shown in figure 2d. The structure shows that gp4 indeed adopts an intramolecular antiparallel coiled-coil topology. The interhelical interface is formed by a hydrophobic core between residues M14, L18, L21 and L25 of α1 and residues W33, Q37 L40 and L44 of α2. The hydrophobic core is stabilized by a salt bridge between residues R17 (α1) and E36 (α2) (Figure 2e) and two hydrogen bonds between the side chain atoms T22-Oγ_1_ and Q36-NHε_22_ and between R11-NHη1 and carboxyl group at the C-terminus. The α2 is tilted from the α1 helix plane, forming an interhelical angle of 30° (Figure 2d). gp4 is a basic peptide (predicted pI=10.5) with a predominantly positive electrostatic surface. The positive charges are more clustered on the surface of α2, comprising six basic amino acids with the side chains exposed to the solvent (Figure 2f).

### Structural study of MvaT-gp4 complex

To uncover the molecular basis of gp4 inhibitory function on MvaT DNA bridging activity, we first investigated the biophysical properties of the MvaT-gp4 complex. It was previously suggested that gp4 interacts with the MvaT oligomerization domain and linker region (residues 1–80) (36). To substantiate this proposal, we first studied the interaction of gp4 with the truncated NTD (residues 1-62) and the DBD (residues 79-124) of MvaT using a magnetic bead pull-down assay (See M&M). The assay shows that gp4 binds to the NTD but not to the truncated DBD (Figure S2b). These results also suggest that the linker region (65-80) may not be required for the formation of the MvaT-gp4 complex.

An isothermal titration calorimetry (ITC) experiment was conducted to determine the stoichiometry and the dissociation constant (K_D_) of the MvaT_2_-gp4 complex. The fit of the titration data yields a K_D_ of 170 ± 14 nM with a number of binding sites N= 0.56 (Figure 3a, See M&M). These results indicate a tight binding between the two proteins with a stoichiometry of 2:1 (MvaT:gp4). The complex appears to be formed by the interaction of one molecule of gp4 with two molecules of MvaT. As the MvaT variant used (F36D/M44D) can only form a dimer through the interaction between the two N-terminal α-helices (site1), we hypothesized that gp4 mainly interacts with site 1, to satisfy the stoichiometry of the complex.

**Figure 3:**
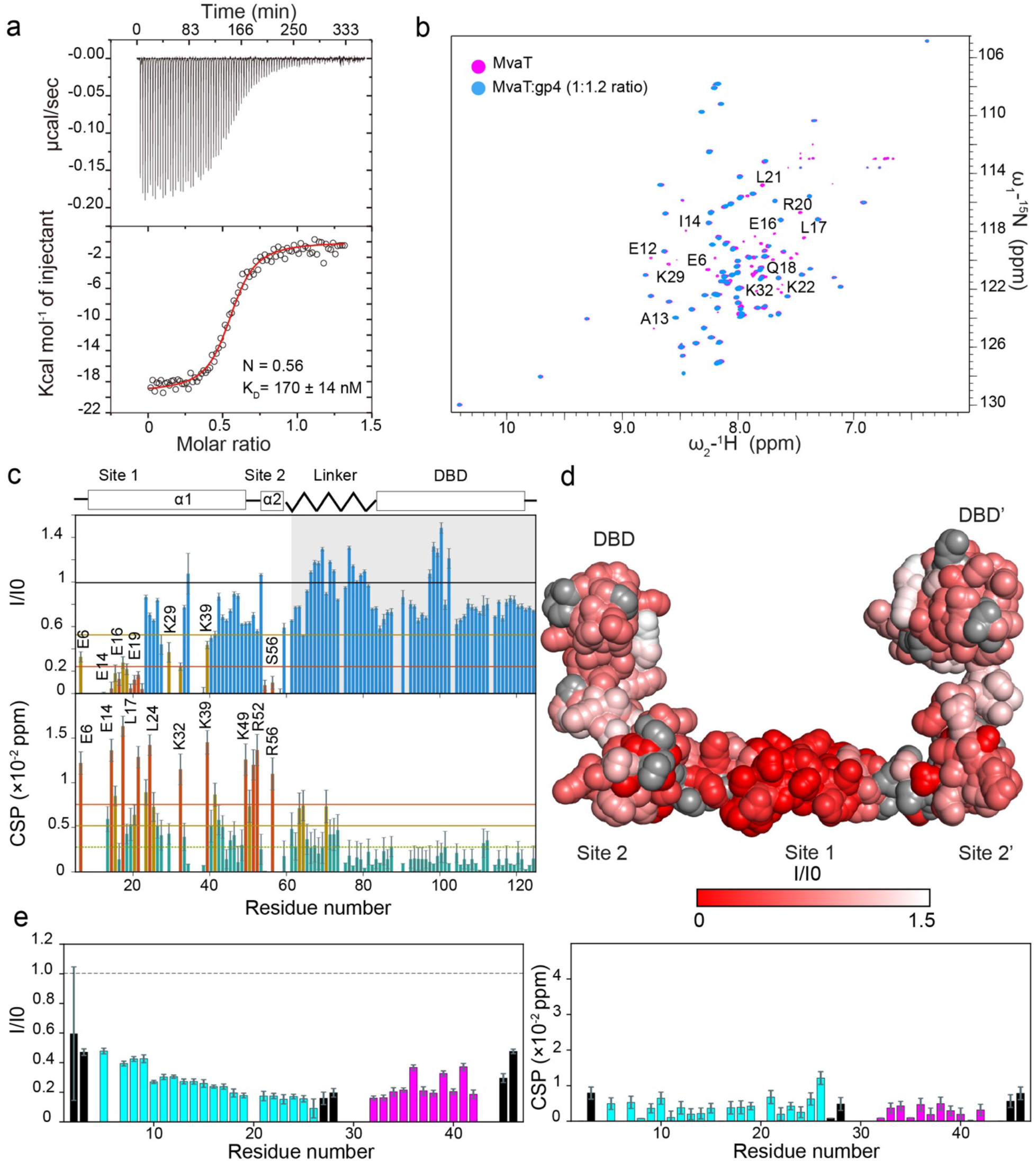
Gp4 interacts with the N-terminal and DBD domains of MvaT. (a) ITC titration of gp4 (230 µM) into MvaT_2_ (monomer concentration = 16 µM) using 2 µl of injection volume. The red line represents the best fit using a one set of sites model (see M&M). The fitting yields a K_D_ = 170 ± 14 nM and N = 0.56. (b) The overlay between ^15^N-^1^H TROSY spectra of the MvaT_2_ in the free state (magenta) and at a molar ratio of MvaT:gp4 of 1:1.2 (blue). Amide resonances which experience severe line broadening upon titration are labelled. (c) The upper panel depicts the ^15^N-^1^H TROSY spectra peak intensities ratio of MvaT in the presence of 1.2 molar ratio of gp4 (I) and the free state (I0) versus the protein residue number. Lower panel shows weighted average CSP of the MvaT resonances between the same points of titration. Resonances with CSP more than two (orange line) or one (yellow line) standard deviation (SD) from the 10% trimmed mean (green dashed line) are labelled and shown in orange, yellow and green bars, respectively. The grey shaded region highlights the changes in the peak intensities of the DBD-linker resonances of MvaT. (d) Mapping of the residues with a reduction in their peak intensities (red/white gradient), upon addition of gp4, on the surface of the MvaT_2_ structural model (18). Residues with no data are colored in grey spheres. (e) Analysis of the NMR titration of ^15^N-gp4 with unlabeled MvaT_2_. The left panel represents the change in the amide resonance peak intensities (I/I0) at a ^15^N-gp4:MvaT 1.2:1 molar ratio. The right panel shows weighted average CSP of the gp4 resonances between the free and bound states. Cyan bars are for amides of gp4 α1, magenta bars are for α2 amides and black bars are for residues of the loop regions.

To corroborate this hypothesis, we performed a NMR titration of a ^15^N labelled MvaT_2_ sample with unlabeled gp4, under the same experimental conditions as used in the ITC experiment. The addition of gp4 to MvaT_2_ induced a decrease in the intensity of multiple resonances in the protein spectrum with minor changes in chemical shift positions of the dimerization site1 residues (Figure 3b,c). At a molar ratio of MvaT:gp4 of 1:1.2, the ratio between the intensities of the peaks of the free MvaT_2_ (I0) and MvaT_2_ bound to gp4 (I) shows that the amide resonances of the dimerization site 1 (residues 1-22) are the most affected upon complex formation (Figure 3c,d). These observations indicate that gp4 binds to the coiled-coil region of the MvaT_2_ dimerization site 1.

A line broadening is also observed for several amide resonances of the DBD (Figure 3c). This suggests a simultaneous interaction of gp4 with the DBD and the NTD within MvaT_2_. These interactions are possibly too weak to be probed by the pull-down assay when the truncated DBD is used. Indeed, NMR titration of the ^15^N gp4 with the DBD does not show significant changes in the peptide HSQC spectrum even at a molar ratio of gp4:DBD of 1:3 (Figure S2c).

Within MvaT_2_, the DBD is tethered to the N-terminal helices by the linker, which increases its local population near the NTD. Therefore, the tethering by the linker might promote the occurrence of intermolecular interactions between the DBD and gp4 when occupying dimerization site 1 of MvaT_2_. The DBD of MvaT includes negatively charged residues on its surface, which may be required for directing the formation of a specific complex with DNA through molecular frustration (18,57). The observed decrease in peak intensities includes these negatively charged patches, which suggests that they interact with the positively exposed surface of gp4.

Previously, we found that at low ionic strength, MvaT_2_ adopts a “half-open” state in which intramolecular electrostatic interactions between the DBD-linker and NTD (site 1) take place. The interface between the two domains includes the positively charged DBD loop 95-ETKGGNHK-102 and the negatively charged fragment 26-QDDKLKKELEDEE-38 of the NTD (18). These intramolecular interactions cause a localized line broadening of the amide resonances of the DBD loop and for residues of the linker region. An increase in ionic strength destabilizes the electrostatic interactions between the two domains and induces an increase in the intensities of the peaks of these MvaT regions (18). Similar to the effects of high ionic strength, an increase in the peak intensities of the loop at 95-102 and linker resonances is observed upon titration with gp4 (Figure 3c,d). This suggests that the binding of gp4 to MvaT_2_ site 1 induces a change in the intramolecular electrostatic interactions and the conformational orientation of the DBD-linker relative to the NTD.

The resonances of the DBD exhibit no significant chemical shift perturbations (CSP) despite the apparent interactions with gp4 (Figure 3c). This observation can possibly be explained by the different binding modes of the DBD to the solvent-exposed surface of gp4. The DBD can also adopt different spatial positions, relative to the NTD, within the conformational ensemble of the MvaT_2_ (open and closed) (18). The average effect of these different chemical environments may result in small CSP for the DBD resonances.

The titration of ^15^N labelled gp4 with unlabeled MvaT_2_ also shows no significant CSP of the amide resonances of gp4 in conjunction with a general line broadening (Figure 3e). Given the high affinity of gp4 and MvaT_2_, the strong reduction in the intensities in the resonances in the NTD of MvaT_2_ upon complex formation, in combination with very small CSP, suggests that binding is in the slow-intermediate regime on the NMR timescale. In the very slow regime new resonances would be expected to appear for the bound state. However, these are not observed, which would be in line with line-broadening caused by slow-intermediate exchange and perhaps binding of gp4 in multiple orientations, similar to an encounter complex stabilized by electrostatic interactions. The general line broadening of the resonances of gp4 upon complex formation is partly due to the increase in the rotational correlation time and may also be caused by the same effect as observed for MvaT_2_. It is clear that the NMR data provide evidence for complex formation and also help to indicate which region on MvaT_2_ is involved in the interactions with gp4.

### Modelling of gp4 inhibition mechanism of MvaT DNA bridging activity

Diffractive crystals of gp4 in complex with the NTD of MvaT wild type (residues 1-62) could not be obtained, possibly because of the high flexibility of the system (see above). Therefore, we used the NMR titration data as experimental restraints to generate structural models of the MvaT_2_-gp4 complex with HADDOCK (51). In our modelling, we only considered the binding of gp4 to the MvaT_2_ NTD (residues 1-22), as it appears to represent the primary interaction site between the two proteins (see M&M). Multiple structures of the MvaT_2_-gp4 dimer complex were obtained (Figure S3a). The lowest energy structure of the complex is shown in Figure 4a. The complex forms a trimeric coiled-coil topology, in which the N-terminal α1 of gp4 is intercalated between the two N-terminal helices of MvaT_2_ site 1. The α2 helix of gp4 exhibits interactions with only one of the MvaT_2_ coiled-coil helices (Figure S3b). The intermolecular interface includes 187 non-bonded contacts mainly formed by electrostatic and hydrophobic interactions between the interfaces of the two proteins. Four salt-bridges stabilize the trimeric coiled-coil located between residues R23, K20, H28 and D36 of α1 and loop region of gp4 with E6, E35 and R20 of site 1 helices. Residues D30 and R43 of gp4 α2 form two additional salt bridges with K31 and E16 of one of the site 1 helices to further stabilize the complex (Figure S3b).

**Figure 4:**
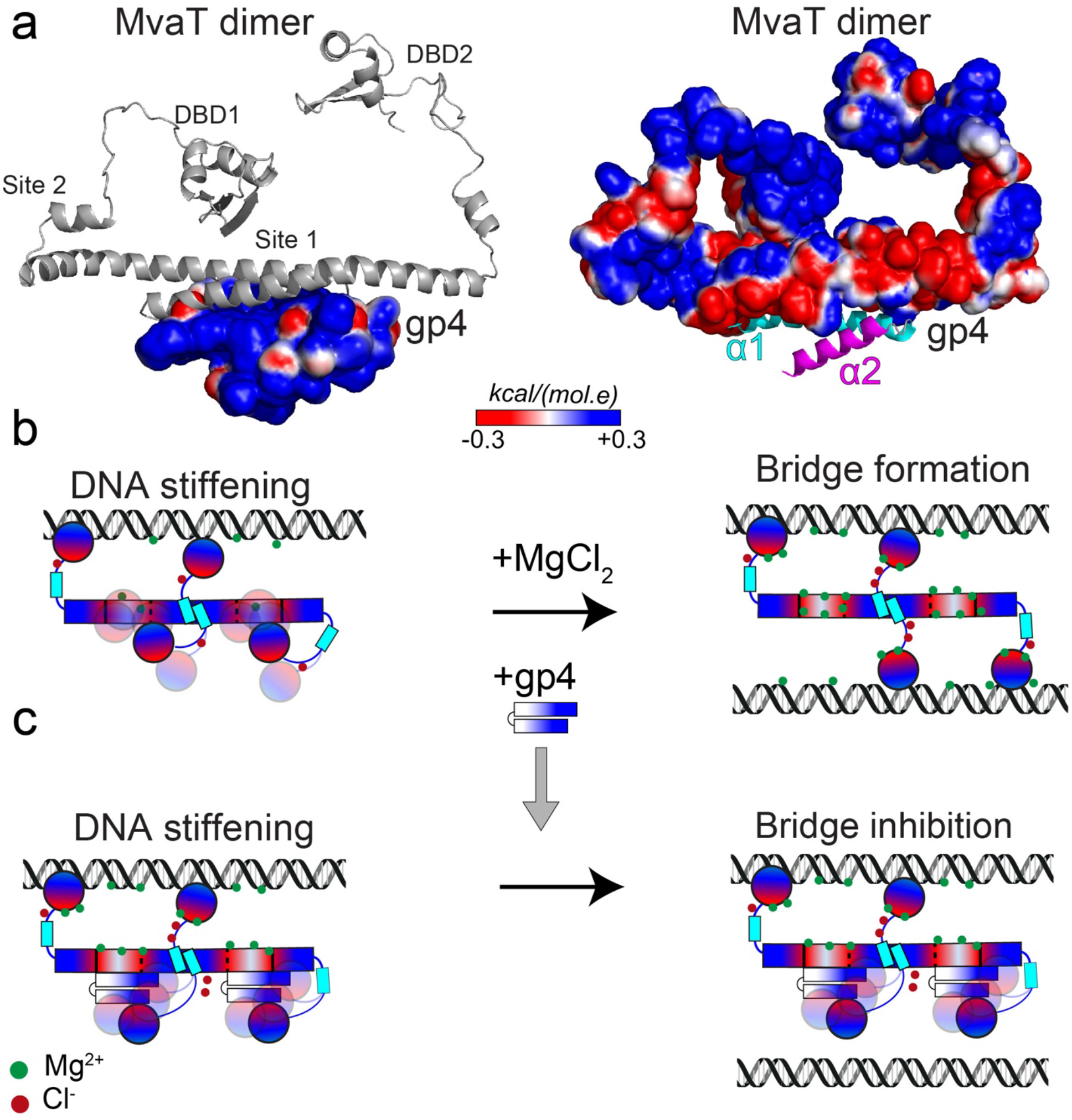
Gp4-MvaT complex and inhibition mechanism of gp4. (a) The best model of the complex between gp4 and MvaT_2_ NTD generated by HADDOCK, using the NMR titration data as restraints, is shown. The electrostatic surfaces of gp4 (right) and MvaT_2_ (left) were generated by CHARMM-GUI (56). (c) The switching mechanism of a MvaT nucleofilament between DNA stiffening and bridging under the influence of salt (18). (c) Schematic representation of the gp4 anti-bridging mechanism. Note that upon interaction of the gp4 with site 1, one of the DBD’s of MvaT_2_ becomes restrained by electrostatic interactions with gp4. The DBD might bind to gp4 on different sites. The electrostatic surfaces of MvaT_2_ and gp4 are shown schematically in a red (negative)/white (neutral)/blue (positive) colour gradient. Green and red dots are for ions (Mg^2+^) and counter ions (Cl^-^), respectively.

Our *in vitro* assays show that in the presence of gp4, the MvaT oligomers are not able to interact with a second DNA duplex to form a bridge. This situation resembles the low ionic strength condition, in which the MvaT oligomers can form filaments yet not bridge DNA. This condition is attributed to the sequestration of one of the DBD’s within the MvaT protomers by electrostatic interactions with the NTD. An increase in ionic strength destabilizes these interdomain interactions and shifts the conformational equilibrium of the MvaT protomer towards an “open state”, able to bridge two DNA duplexes (Figure 4b) (18). Although our bridging assay was performed at high ionic strength (20 mM MgCl_2_), MvaT remains unable to bridge two DNA duplexes in the presence of gp4 (see M&M).

Our structural studies suggest that the primary binding site of gp4 is on the dimerization site 1 of MvaT protomers. Upon complex formation, a structural rearrangement of the MvaT protomer seems to occur, in which one of the DBD’s interacts with the surface of gp4 that is not involved in the interface with MvaT_2_ site 1 (Figure 4c). These interactions appear to be mediated through electrostatic forces. As a result, gp4 will act as an “electostatic zipper” between the NTD and DBD of the MvaT protomers and stabilize a structure resembling the “half-open” conformational state within the protein oligomers (Figure 4c). Consequently, the MvaT-gp4 nucleofilaments will exhibit weak DNA bridging activity, despite forming high-order oligomers along double-stranded DNA.

### Validation of the molecular mechanism of MvaT interference by gp4

To validate the proposed molecular model for the mechanism of MvaT inhibition by gp4, we generated a gp4 variant in which the solvent-exposed residue K42 and K45 of the α2 helix are substituted with glutamic acids (Figure 4a). These charge inversion mutations were aimed introducing repulsive electrostatic forces between gp4 and the DBD to shift the conformational equilibrium of the MvaT protomers toward an “open state”, thus restoring their DNA bridging capacity. The anti-bridging activity of gp4 K42E/K45E was monitored by the pull-down bridging using gp4 WT as a reference. Indeed, a 50 to 30% loss of gp4 inhibitory function is observed at 0.9 and 1.2 µM gp4 respectively, added to 2.4 µM MvaT (Figure 5b). However, at 1.2 µM gp4 started to aggregate on the magnetic beads, making measurement at higher molar ratio impossible. The stoichiometry of the complex appears equal to 0.52, similar to that obtained for the MvaT_2_ titration with gp4 WT, as determined by ITC (Figure 5c). However, a tenfold increase in the K_D_ value is observed (1.36 ± 0.06 μM), possibly due to loss of some intermolecular interactions within the complex. To verify this hypothesis, we performed a NMR titration of ^15^N labelled MvaT_2_ with the mutated gp4. The gp4 mutant binds to the dimerization site 1 as concluded from the observed CSP and the reduction in the ratio of peak intensities (Figure 5d). In contrast to the titration with WT gp4, no line broadening is observed for the DBD-linker resonances. These results suggest a loss of the intermolecular interaction between gp4 K42E/K45E and the DBD-linker of the MvaT_2_. Collectively, these data support our model in which gp4 is acting as an electrostatic stabilizer of a structure resembling the “half-open” state of the MvaT protomers for the selective inhibition of their DNA bridging activity.

**Figure 5:**
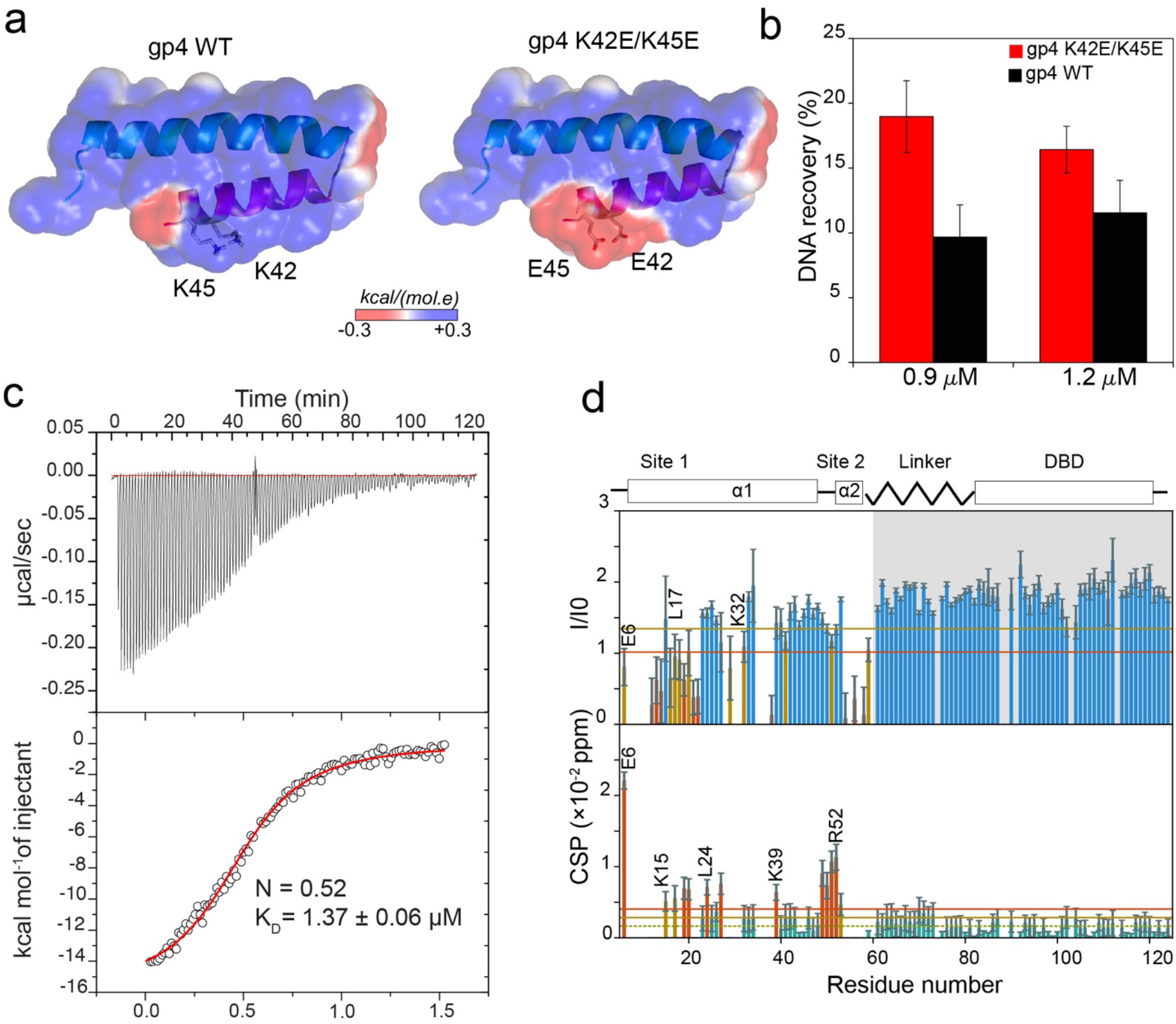
Validation of gp4 anti-bridging mechanism by point mutagenesis: (a) electrostatic surface of gp4 WT and gp4 K42E/K45 mutant. (b) Inhibitory effects of gp4 mutant (red bars) and gp4 (black bars) on MvaT DNA bridging activity at 0.9 and 1.2 μM of gp4 using a constant concentration of MvaT of 2.4 μM (see M&M). (c) ITC of MvaT_2_ with gp K42E/K45E. The red line shows the best fit of the data. (d) Analysis of the NMR titration data of ^15^N MvaT_2_ with gp4 K42E/K45E. The upper panel depicts the ^15^N-^1^H TROSY spectra peak intensities ratio of MvaT_2_ in the presence of 1.2 molar ratio of gp4 (I) and the free state (I0) versus the protein residue number. Note the two-fold increase in the peak intensities of the DBD-linker region (shaded in grey) due to the loss of the intramolecular electrostatic interactions between the DBD-linker -NTD within MvaT_2_ subunits(18) and the intermolecular interactions with gp4. Lower panel shows weighted average CSP of the MvaT resonances between the same points of titration. Resonances with CSP more than two (orange line) or one (yellow line) standard deviation (SD) from the 10% trimmed mean (green dashed line) are labelled and shown in orange, yellow and green bars, respectively.

## Discussion

Bacterial species and their associated bacteriophages are in a relentless evolutionary arms race in which they develop defense and counter-defense mechanisms to assure survival. One of the mechanisms that phages deploy is the production of “early proteins” to shut down host defense mechanisms and co-opt host-cell machinery for efficient infection (58,59). Despite representing a rich source of antimicrobial molecules, most of these phage early proteins are considered nonessential, as they are not required for phage production. Therefore, the structure, target, and mode of actions of these proteins have remained vastly unexplored.

One of the first line of Enteric bacteria resistance systems is the silencing of foreign (viral) DNA by H-NS family proteins. However, bacteriophages encode proteins that counteract this host-defense system. In this work, we have investigated the molecular mechanism by which the phage LUZ24 “early protein” gp4 functionally modulates the *P. aeruginosa* H-NS family protein MvaT to markedly inhibit the growth of this important pathogen. Using (single-molecule) biophysical methods, we found that gp4 specifically targets the formation and maintenance of bridged DNA complexes by MvaT without interfering with MvaT oligomerization along DNA. This situation resembles the low ionic strength conditions under which the MvaT protomers adopt a “half-open” conformational state, incompetent to bridge two DNA duplexes while forming lateral filaments along DNA. This structural configuration results from intramolecular electrostatic interactions between the oppositely charged DBD and dimerization site 1 of the MvaT protomers (18). Structural studies revealed that gp4 simultaneously interacts with dimerization site 1 and the DBD of the MvaT protomers to stabilize a mimic of the “half-open” state through electrostatic interactions. Introduction of repulsive electrostatic forces at the interface of gp4 and the DBD of the MvaT dimer results in diminished anti-bridging activity of gp4.

Bacteria encode several proteins that modulate the structure and function of H-NS-DNA complex. For instance, truncated H-NS (H-NST) derivatives, which lack the C-terminal DNA binding domain, were found in large genomic islands of pathogenic *E. coli* strains (60). It was suggested that H-NST family members act as antagonists of H-NS function by preventing its correct oligomerization through heterodimerization with the N-terminal dimerization domain of H-NS. The expression of YdgT stimulates DNA bridging by enhancing the cooperativity of the lateral nucleofilament formation of H-NS (19). Binding of Hha/YdgT family proteins to the H-NS dimerization site (residues 1-45) stimulates H-NS DNA bridging activity (19), and thus enhances transcriptional silencing of foreign genes in bacteria (61). Hha forms electrostatic complexes with the “hand-shake” fold of H-NS dimerization site 1 with a 1:1 stoichiometry (Hha: H-NS monomer) and exposes a positively charged surface (62,63). It was proposed that the exposed surface of Hha enhances the binding affinity of H-NS filament to DNA and stabilizes the bridging complex to stimulate pausing of RNA polymerase (64). The LUZ24 gp4 seems to operate analogously to Hha regarding the electrostatic interactions with dimerization site 1 of H-NS family proteins and the exposure of a positively charged surface. Nevertheless, the stoichiometry, topology and orientation of the complex appear to differ because of the deviations in the fold architectures of MvaT and H-NS site 1 (coiled-coil vs. hand-shake) and of gp4 and Hha, which result in opposite effects on the bridging activity of the two H-NS family members.

DNA bridging by H-NS proteins is thought to be crucial for gene silencing and for the global organization of Enteric bacteria genomes (65-67). Additionally, molecular studies have suggested that a switch between linear and bridging nucleofilaments of H-NS family proteins drives the bacterial adaptation to environmental factors (18,19). The selective inhibition of MvaT DNA bridging activity by gp4, which translates into an adverse impact on growth of *P. aeruginosa* provides additional evidence of the importance of DNA bridging by H-NS family proteins in maintaining bacterial homeostasis. *P. aeruginosa* also encodes MvaU, a paralog of MvaT that binds to many of the same chromosomal regions and also bridges DNA duplexes (15,68). Nevertheless, despite the structural and functional similarity between these two proteins, gp4 appears to target MvaT selectively and requires its presence for cytotoxicity. Future studies on the impact of gp4 on nucleoid organization and regulation of gene expression in *P. aeruginosa* might provide more insight into the antimicrobial mechanism of this phage early protein.

In conclusion, a better understanding of how viral proteins counteract H-NS proteins is essential for designing molecules that modulate phage infections. Potential applications of such molecules can be envisioned in biotechnology, where phage infections are a threat to large scale culturing, or in phage therapy where they may enhance the ability to infect and kill their host. Moreover, due to their global role in gene regulation in bacteria, H-NS family proteins are potential drug targets to mitigate recalcitrant bacterial infections. A mechanistic understanding at the molecular level of how H-NS family protein activity is modulated will prove key to such developments.

## Data availability

The datasets generated during the current study are available from the 4TU repository (https://data.4tu.nl) under DOI: https://doi.org/10.4121/14207180. The resonance assignments of gp4 were deposited in the BMRB data bank under the accession number 28112. The NMR structures of gp4 were deposited in the PDB bank under the accession code 6YSE.

## Acknowledgments/funding

This research was supported by grants from the Netherlands Organization for Scientific Research [VICI 016.160.613] (R.T.D.), the Human Frontier Science Program (HFSP) [RGP0014/2014] (R.T.D.), a grant from the China Scholarship Council (CSC) [201506880001] (L.Q.), and National Institutes of Health grant AI105013 (S.L.D). F.B.B thanks the Leiden University-Gratama fund [2020-09/W20386-1-GSL].

## Contributions

F.B.B., L.Q., A.N.V., A.M.L., A.M.E., N.B., A.B., performed the experiments. F.B.B., L.Q., A.N.V.,

A.M.L., A.M.E., N.B., A.L.B., M.U., S.L.D. and R.T.D contributed to data analysis and discussion. F.B.B. and R.T.D. supervised the project. F.B.B. and R.T.D. wrote the manuscript. All authors reviewed and corrected the manuscript.

## References

1. Bergh, Ø., Børsheim, K.Y., Bratbak, G. and Heldal, M. (1989) High abundance of viruses found in aquatic environments. Nature, 340, 467–468.

2. Wommack, K.E. and Colwell, R.R. (2000) Virioplankton: viruses in aquatic ecosystems. Microbiology and molecular biology reviews, 64, 69–114.

3. Stern, A. and Sorek, R. (2011) The phage-host arms race: shaping the evolution of microbes. Bioessays, 33, 43–51.

4. Chibani-Chennoufi, S., Bruttin, A., Dillmann, M.-L. and Brüssow, H. (2004) Phage-host interaction: an ecological perspective. Journal of bacteriology, 186, 3677–3686.

5. Stone, E., Campbell, K., Grant, I. and McAuliffe, O. (2019) Understanding and exploiting phage–host interactions. Viruses, 11, 567.

6. Enikeeva, F.N., Severinov, K.V. and Gelfand, M.S. (2010) Restriction–modification systems and bacteriophage invasion: Who wins? Journal of theoretical biology, 266, 550–559.

7. Dupuis, M.-É., Villion, M., Magadán, A.H. and Moineau, S. (2013) CRISPR-Cas and restriction–modification systems are compatible and increase phage resistance. Nature communications, 4, 1–7.

8. Pfeifer, E., Hünnefeld, M., Popa, O. and Frunzke, J. (2019) Impact of xenogeneic silencing on phage–host interactions. Journal of molecular biology, 431, 4670–4683.

9. Flores-Ríos, R., Quatrini, R. and Loyola, A. (2019) Endogenous and foreign nucleoid-associated proteins of bacteria: occurrence, interactions and effects on mobile genetic elements and host’s biology. Computational and structural biotechnology journal, 17, 746–756.

10. Navarre, W. (2016) The impact of gene silencing on horizontal gene transfer and bacterial evolution. Advances in microbial physiology, 69, 157–186.

11. Hüttener, M., Paytubi, S. and Juárez, A. (2015) Success in incorporating horizontally transferred genes: the H-NS protein. Trends in microbiology, 23, 67–69.

12. Singh, K., Milstein, J.N. and Navarre, W.W. (2016) Xenogeneic silencing and its impact on bacterial genomes. Annual review of microbiology, 70, 199–213.

13. Qin, L., Erkelens, A., Ben Bdira, F. and Dame, R. (2019) The architects of bacterial DNA bridges: a structurally and functionally conserved family of proteins. Open biology, 9, 190223.

14. Navarre, W.W., McClelland, M., Libby, S.J. and Fang, F.C. (2007) Silencing of xenogeneic DNA by H-NS—facilitation of lateral gene transfer in bacteria by a defense system that recognizes foreign DNA. Genes & Development, 21, 1456–1471.

15. Castang, S., McManus, H.R., Turner, K.H. and Dove, S.L. (2008) H-NS family members function coordinately in an opportunistic pathogen. Proceedings of the National Academy of Sciences, 105, 18947–18952.

16. Gordon, B.R., Imperial, R., Wang, L., Navarre, W.W. and Liu, J. (2008) Lsr2 of Mycobacterium represents a novel class of H-NS-like proteins. Journal of bacteriology, 190, 7052–7059.

17. Smits, W.K. and Grossman, A.D. (2010) The transcriptional regulator Rok binds A+ T-rich DNA and is involved in repression of a mobile genetic element in Bacillus subtilis. PLoS Genet, 6, e1001207.

18. Qin, L., Bdira, F.B., Sterckx, Y.G., Volkov, A.N., Vreede, J., Giachin, G., van Schaik, P., Ubbink, M. and Dame, R.T. (2020) Structural basis for osmotic regulation of the DNA binding properties of H-NS proteins. Nucleic acids research, 48, 2156–2172.

19. van der Valk, R.A., Vreede, J., Qin, L., Moolenaar, G.F., Hofmann, A., Goosen, N. and Dame, R.T. (2017) Mechanism of environmentally driven conformational changes that modulate H-NS DNA-bridging activity. Elife, 6, e27369.

20. Winardhi, R.S., Fu, W., Castang, S., Li, Y., Dove, S.L. and Yan, J. (2012) Higher order oligomerization is required for H-NS family member MvaT to form gene-silencing nucleoprotein filament. Nucleic acids research, 40, 8942–8952.

21. Qu, Y., Lim, C.J., Whang, Y.R., Liu, J. and Yan, J. (2013) Mechanism of DNA organization by Mycobacterium tuberculosis protein Lsr2. Nucleic acids research, 41, 5263–5272.

22. Dame, R.T., Wyman, C. and Goosen, N. (2000) H-NS mediated compaction of DNA visualised by atomic force microscopy. Nucleic acids research, 28, 3504–3510.

23. Dame, R.T., Noom, M.C. and Wuite, G.J. (2006) Bacterial chromatin organization by H-NS protein unravelled using dual DNA manipulation. Nature, 444, 387–390.

24. Dillon, S.C. and Dorman, C.J. (2010) Bacterial nucleoid-associated proteins, nucleoid structure and gene expression. Nature Reviews Microbiology, 8, 185–195.

25. Dorman, C.J. (2013) Genome architecture and global gene regulation in bacteria: making progress towards a unified model? Nature Reviews Microbiology, 11, 349–355.

26. Dame, R.T. (2005) The role of nucleoid-associated proteins in the organization and compaction of bacterial chromatin. Molecular microbiology, 56, 858–870.

27. Dame, R.T., Rashid, F.-Z.M. and Grainger, D.C. (2020) Chromosome organization in bacteria: mechanistic insights into genome structure and function. Nature Reviews Genetics, 21, 227–242.

28. Liu, Q. and Richardson, C.C. (1993) Gene 5.5 protein of bacteriophage T7 inhibits the nucleoid protein H-NS of Escherichia coli. Proceedings of the National Academy of Sciences, 90, 1761–1765.

29. Ali, S.S., Beckett, E., Bae, S.J. and Navarre, W.W. (2011) The 5.5 protein of phage T7 inhibits H-NS through interactions with the central oligomerization domain. Journal of bacteriology, 193, 4881–4892.

30. Zhu, B., Lee, S.-J., Tan, M., Wang, E.-D. and Richardson, C.C. (2012) Gene 5.5 protein of bacteriophage T7 in complex with Escherichia coli nucleoid protein H-NS and transfer RNA masks transfer RNA priming in T7 DNA replication. Proceedings of the National Academy of Sciences, 109, 8050–8055.

31. Ho, C.-H., Wang, H.-C., Ko, T.-P., Chang, Y.-C. and Wang, A.H.-J. (2014) The T4 phage DNA mimic protein Arn inhibits the DNA binding activity of the bacterial histone-like protein H-NS. Journal of Biological Chemistry, 289, 27046–27054.

32. Patterson-West, J., Tai, C., Son, B., Hsieh, M., Iben, J. and Hinton, D. (2021). s Note: MDPI stays neu-tral with regard to jurisdictional clai-ms in ….

33. Yasunobu, K., Kayoko, Y., Hiromitu, T. and Fumio, I. (1993) Propagation of phage Mu in IHF-dencient Escherichia coli in the absence of the H-NS histone-like protein. Gene, 126, 93–97.

34. van Ulsen, P., Hillebrand, M., Zulianello, L., van de Putte, P. and Goosen, N. (1996) Integration host factor alleviates the H-NS-mediated repression of the early promoter of bacteriophage Mu. Molecular microbiology, 21, 567–578.

35. Skennerton, C.T., Angly, F.E., Breitbart, M., Bragg, L., He, S., McMahon, K.D., Hugenholtz, P. and Tyson, G.W. (2011) Phage encoded H-NS: a potential achilles heel in the bacterial defence system. PloS one, 6, e20095.

36. Wagemans, J., Delattre, A.-S., Uytterhoeven, B., De Smet, J., Cenens, W., Aertsen, A., Ceyssens, P.-J. and Lavigne, R. (2015) Antibacterial phage ORFans of Pseudomonas aeruginosa phage LUZ24 reveal a novel MvaT inhibiting protein. Frontiers in microbiology, 6, 1242.

37. Lippa, A.M., Gebhardt, M.J. and Dove, S.L. (2020) H-NS-like proteins in Pseudomonas aeruginosa coordinately silence intragenic transcription. Molecular Microbiology.

38. Gibson, D.G., Young, L., Chuang, R.-Y., Venter, J.C., Hutchison, C.A. and Smith, H.O. (2009) Enzymatic assembly of DNA molecules up to several hundred kilobases. Nature methods, 6, 343–345.

39. van der Valk, R.A., Qin, L., Moolenaar, G.F. and Dame, R.T. (2018), Bacterial Chromatin. Springer, pp. 199–209.

40. Vallet-Gely, I., Donovan, K.E., Fang, R., Joung, J.K. and Dove, S.L. (2005) Repression of phase-variable cup gene expression by H-NS-like proteins in Pseudomonas aeruginosa. Proceedings of the National Academy of Sciences, 102, 11082–11087.

41. Hmelo, L.R., Borlee, B.R., Almblad, H., Love, M.E., Randall, T.E., Tseng, B.S., Lin, C., Irie, Y., Storek, K.M. and Yang, J.J. (2015) Precision-engineering the Pseudomonas aeruginosa genome with two-step allelic exchange. Nature protocols, 10, 1820.

42. Meisner, J. and Goldberg, J.B. (2016) The Escherichia coli rhaSR-PrhaBAD inducible promoter system allows tightly controlled gene expression over a wide range in Pseudomonas aeruginosa. Applied and environmental microbiology, 82, 6715–6727.

43. Hoang, T.T., Karkhoff-Schweizer, R.R., Kutchma, A.J. and Schweizer, H.P. (1998) A broad-host-range Flp-FRT recombination system for site-specific excision of chromosomally-located DNA sequences: application for isolation of unmarked Pseudomonas aeruginosa mutants. Gene, 212, 77–86.

44. Lee, W., Tonelli, M. and Markley, J.L. (2015) NMRFAM-SPARKY: enhanced software for biomolecular NMR spectroscopy. Bioinformatics, 31, 1325–1327.

45. Delaglio, F., Grzesiek, S., Vuister, G.W., Zhu, G., Pfeifer, J. and Bax, A. (1995) NMRPipe: a multidimensional spectral processing system based on UNIX pipes. Journal of biomolecular NMR, 6, 277–293.

46. Vranken, W.F., Boucher, W., Stevens, T.J., Fogh, R.H., Pajon, A., Llinas, M., Ulrich, E.L., Markley, J.L., Ionides, J. and Laue, E.D. (2005) The CCPN data model for NMR spectroscopy: development of a software pipeline. Proteins, 59, 687–696.

47. Cheung, M.S., Maguire, M.L., Stevens, T.J. and Broadhurst, R.W. (2010) DANGLE: A Bayesian inferential method for predicting protein backbone dihedral angles and secondary structure. Journal of magnetic resonance, 202, 223–233.

48. Guntert, P. and Buchner, L. (2015) Combined automated NOE assignment and structure calculation with CYANA. Journal of biomolecular NMR, 62, 453–471.

49. Brunger, A.T., Adams, P.D., Clore, G.M., DeLano, W.L., Gros, P., Grosse-Kunstleve, R.W., Jiang, J.S., Kuszewski, J., Nilges, M., Pannu, N.S. et al. (1998) Crystallography & NMR system: A new software suite for macromolecular structure determination. Acta crystallographica. Section D, Biological crystallography, 54, 905–921.

50. Schwieters, C.D., Kuszewski, J.J., Tjandra, N. and Clore, G.M. (2003) The Xplor-NIH NMR molecular structure determination package. Journal of magnetic resonance, 160, 65–73.

51. Van Zundert, G., Rodrigues, J., Trellet, M., Schmitz, C., Kastritis, P., Karaca, E., Melquiond, A., van Dijk, M., De Vries, S. and Bonvin, A. (2016) The HADDOCK2. 2 web server: user-friendly integrative modeling of biomolecular complexes. Journal of molecular biology, 428, 720–725.

52. Castang, S. and Dove, S.L. (2010) High-order oligomerization is required for the function of the H-NS family member MvaT in Pseudomonas aeruginosa. Molecular microbiology, 78, 916–931.

53. Dame, R.T., Luijsterburg, M.S., Krin, E., Bertin, P.N., Wagner, R. and Wuite, G.J. (2005) DNA bridging: a property shared among H-NS-like proteins. Journal of Bacteriology, 187, 1845–1848.

54. Myszka, D.G. and Chaiken, I.M. (1994) Design and characterization of an intramolecular antiparallel coiled coil peptide. Biochemistry, 33, 2363–2372.

55. Drozdetskiy, A., Cole, C., Procter, J. and Barton, G.J. (2015) JPred4: a protein secondary structure prediction server. Nucleic acids research, 43, W389–W394.

56. Jo, S., Kim, T., Iyer, V.G. and Im, W. (2008) CHARMM-GUI: a web-based graphical user interface for CHARMM. Journal of computational chemistry, 29, 1859–1865.

57. Marcovitz, A. and Levy, Y. (2011) Frustration in protein–DNA binding influences conformational switching and target search kinetics. Proceedings of the National Academy of Sciences, 108, 17957–17962.

58. De Smet, J., Hendrix, H., Blasdel, B.G., Danis-Wlodarczyk, K. and Lavigne, R. (2017) Pseudomonas predators: understanding and exploiting phage–host interactions. Nature Reviews Microbiology, 15, 517.

59. Roucourt, B. and Lavigne, R. (2009) The role of interactions between phage and bacterial proteins within the infected cell: a diverse and puzzling interactome. Environmental microbiology, 11, 2789–2805.

60. Williamson, H.S. and Free, A. (2005) A truncated H-NS-like protein from enteropathogenic Escherichia coli acts as an H-NS antagonist. Molecular microbiology, 55, 808–827.

61. Ueda, T., Takahashi, H., Uyar, E., Ishikawa, S., Ogasawara, N. and Oshima, T. (2013) Functions of the Hha and YdgT proteins in transcriptional silencing by the nucleoid proteins, H-NS and StpA, in Escherichia coli. DNA research, 20, 263–271.

62. Ali, S.S., Whitney, J.C., Stevenson, J., Robinson, H., Howell, P.L. and Navarre, W.W. (2013) Structural insights into the regulation of foreign genes in Salmonella by the Hha/H-NS complex. Journal of Biological Chemistry, 288, 13356–13369.

63. Cordeiro, T.N., García, J., Bernadó, P., Millet, O. and Pons, M. (2015) A three-protein charge zipper stabilizes a complex modulating bacterial gene silencing. Journal of Biological Chemistry, 290, 21200–21212.

64. Boudreau, B.A., Hron, D.R., Qin, L., Van Der Valk, R.A., Kotlajich, M.V., Dame, R.T. and Landick, R. (2018) StpA and Hha stimulate pausing by RNA polymerase by promoting DNA–DNA bridging of H-NS filaments. Nucleic acids research, 46, 5525–5546.

65. Kotlajich, M.V., Hron, D.R., Boudreau, B.A., Sun, Z., Lyubchenko, Y.L. and Landick, R. (2015) Bridged filaments of histone-like nucleoid structuring protein pause RNA polymerase and aid termination in bacteria. Elife, 4, e04970.

66. Van Der Valk, R.A., Vreede, J., Crémazy, F. and Dame, R.T. (2014) Genomic looping: a key principle of chromatin organization. Journal of molecular microbiology and biotechnology, 24, 344–359.

67. Stoebel, D.M., Free, A. and Dorman, C.J. (2008) Anti-silencing: overcoming H-NS-mediated repression of transcription in Gram-negative enteric bacteria. Microbiology, 154, 2533–2545.

68. Winardhi, R.S., Castang, S., Dove, S.L. and Yan, J. (2014) Single-molecule study on histone-like nucleoid-structuring protein (H-NS) paralogue in Pseudomonas aeruginosa: MvaU bears DNA organization mode similarities to MvaT. PLoS One, 9, e112246.

