## Supplemental material for "Novel anti-repression mechanism of H-NS proteins by a phage “early protein”"

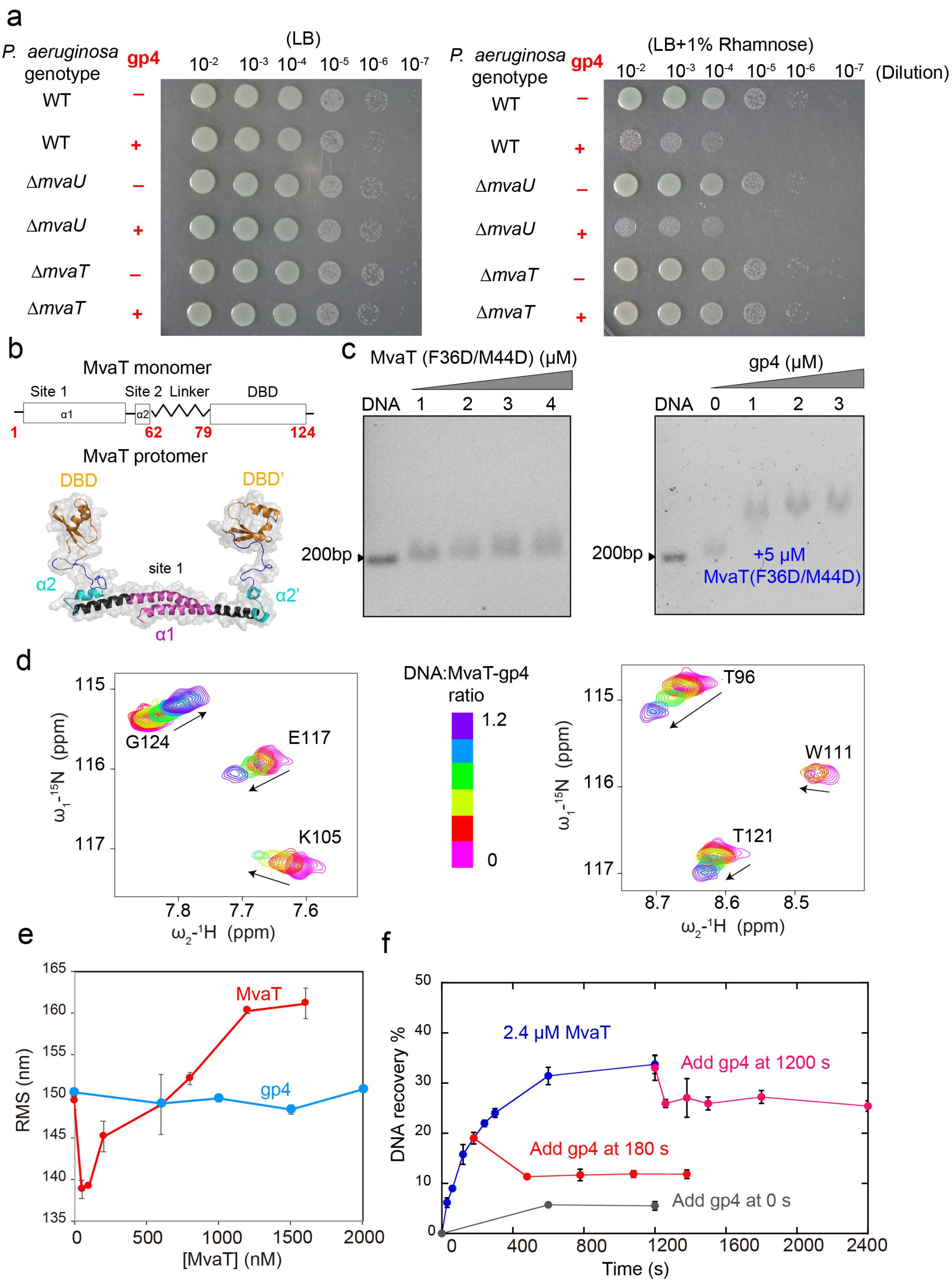


**Figure S1**: (a) Heterologous expression of gp4 *in Pseudomonas aeruginosa* wild-type cells and cells lacking either *mvaU* or *mvaT*. The left panel shows serial dilutions of cells growth in the absence of inducer and the right panel is in the presence of Rhamnose for induction of gp4 expression. (b) MvaT monomer and protomer fold topology. In the lower panel is the structural model of the MvaT protomer adopted from (1). (c) Left panel is for electrophoretic mobility shift assay (EMSA) of a 200bp DNA substrate at different MvaT F36D/M44D concentrations. Right panel is for an EMSA showing DNA binding by 5 μM MvaT F36D/M44D titrated with different concentrations of gp4. (d) NMR titration of $^{15}N$ MvaT F36D/M44D:gp4 complex (1:1.2 molar ratio) with a 20 bp DNA substrate. Representative chemical shift perturbations of the MvaT DBD are shown and indicated by black arrows. (e) The red curve represents the RMS of DNA bound by MvaT at concentrations from 0 to 1600 nM measured by the Tethered Particle Motion in the absence of Mip. The blue curve is for the RMS of DNA titrated by increased concentrations of gp4 between 0 and 2000 nM, in the absence of MvaT. The error bars represent the standard deviation of at least two independent measurements; some error bars are hidden behind the data points. (f) Time dependent bridging assay of DNA-MvaT-DNA complexes. The blue curve represents the kinetics of MvaT-DNA bridge formation without adding gp4. The red and pink curve represent the kinetics of MvaT-DNA bridge formation with Mip introduced at 180 s and 1200 s, respectively. The grey curve represents the introduction of gp4 at the beginning of the assay (t = 0 s). Error bars represent the standard deviation from at least two independent measurements.

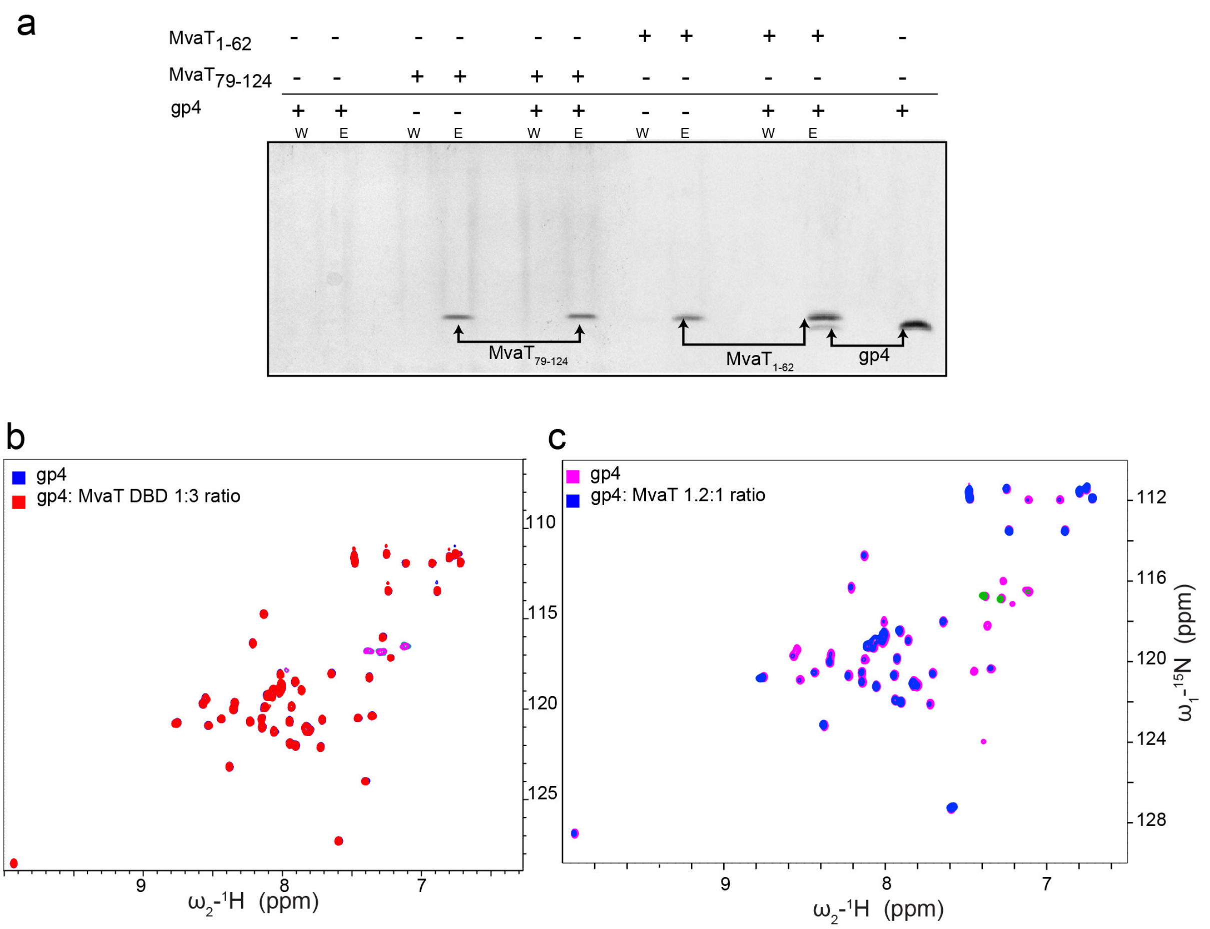


**Figure S2:** (a) His-tag pull down assay of MvaT truncated domains (NTD is 1-62 and DBD is 79-124) in the presence of gp4 analyzed by Tricine gel electrophoresis. W and E are for the flow through of the washing and elution steps, respectively (see M&M). (b) Overlay between $^{15}N$ gp4 HSQC spectra in the absence (blue) and presence of unlabeled MvaT DBD at a gp4:DBD 1:3 molar ratio. Magenta and green are for the folded peaks of gp4 arginine side chains. (c) Overlay between gp4 HSQC spectra in the absence (magenta) and presence (blue) of $MvaT_2$ at a gp4:MvaT 1.2:1 ratio. Light green is for the folded peaks of gp4 arginine side chains.

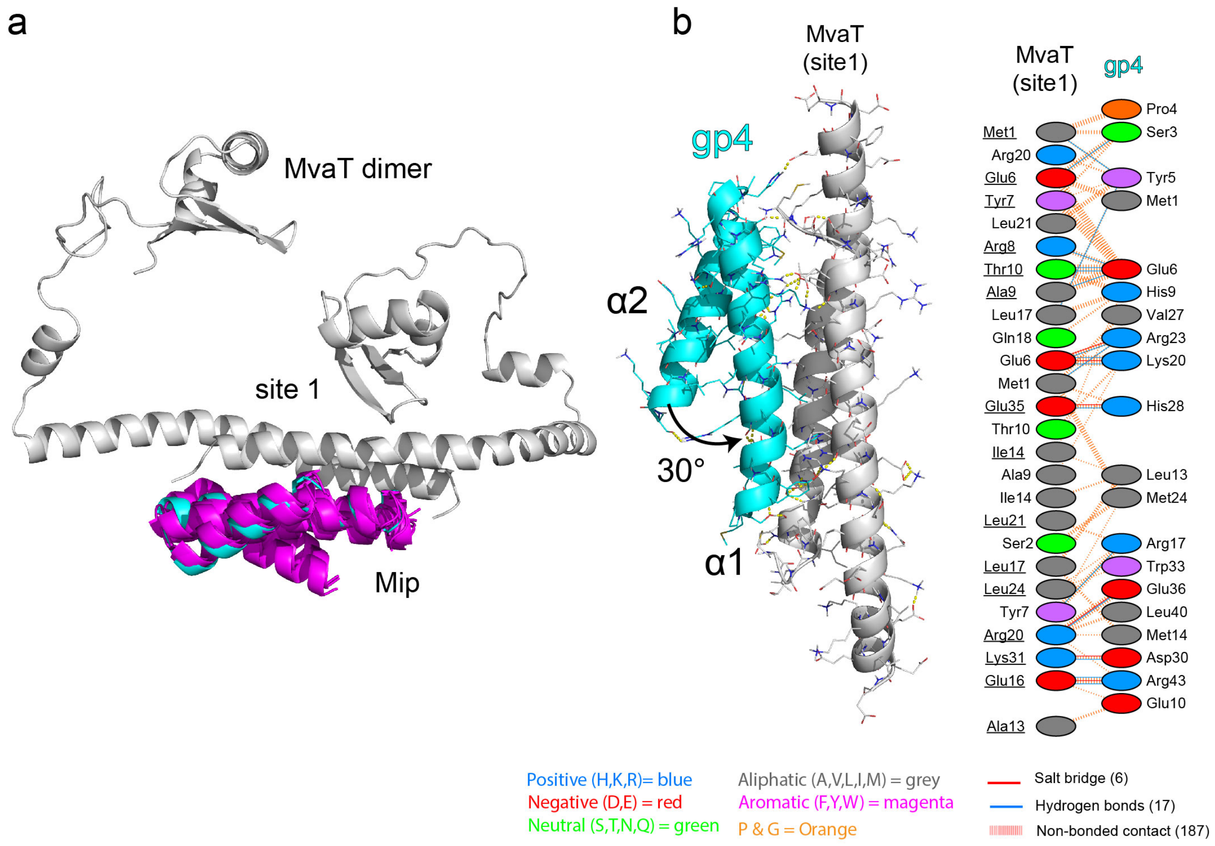


**Figure S3: (a) HADDOCK best cluster of gp4-MvaT dimer complex**. The different orientations of gp4 are shown in magenta cartoon and the MvaT dimer in grey cartoon. The lowest energy structure of gp4 is shown is cyan cartoon. (b) The trimeric coiled-coil complex between gp4 α1 and the two helices of MvaT site 1 is depicted in cartoon. The angle between gp4 helices is indicated. The right panel is for the analysis of the intermolecular interactions within the trimeric coiled-coil by PDBsum (2). The underlined amino acids belong to the second helix of the MvaT dimerization site1. The type and number of the intermolecular interaction are indicated in different colors. The amino acids residues of the complex interface are colored based on the chemical characteristics of their side chains, as indicated.

**Table S1.** Structural statistics over 10 lowest-energy NMR structures of gp4. [a]Root-mean-square deviation from the lowest-energy structure calculated for residues 4-45. [b]Calculated with MolProbity.

| | |
|---|---|
| Distance restraints | |
| Total | 1290 |
| Short-range, $\|i - j\| \leq 1$ | 665 |
| Medium-range, $1 < \|i - j\| < 5$ | 406 |
| Long-range, $\|i - j\| \geq 5$ | 219 |
| Dihedral angle restraints | |
| φ and ψ | 78 |
| Restraints violations | |
| NOE, > 0.5 Å | 0 |
| Dihedral angle, > 5° | 0 |
| Coordinate RMSD,[a] Å | |
| Backbone | 0.32 ± 0.08 |
| Heavy atoms | 1.09 ± 0.16 |
| Ramachandran statistics,[b] % | |
| Favoured | 99.3 |
| Allowed | 0.7 |
| Outliers | 0.0 |

**Table S2.** Statistics of the top cluster of HADDOCK docking.

| **Cluster 3** | |
|---|---|
| HADDOCK score | -103.3 +/- 14.3 |
| Cluster size | 11 |
| RMSD from the overall lowest-energy structure | 1.0 +/- 0.6 |
| Van der Waals energy | 67.0 +/- 7.6 |
| Electrostatic energy | -360.2 +/- 63.5 |
| Desolvation energy | -5.0 +/- 7.0 |
| Restraints violation energy | 407.9 +/- 78.85 |
| Buried Surface Area | 2369.2 +/- 90.2 |
| Z-Score | -2.0 |

**References:**


1. Qin, L., Bdira, F.B., Sterckx, Y.G., Volkov, A.N., Vreede, J., Giachin, G., van Schaik, P., Ubbink, M. and Dame, R.T. (2020) Structural basis for osmotic regulation of the DNA binding properties of H-NS proteins. *Nucleic acids research*, **48**, 2156-2172.
2. Laskowski, R.A., Jabłońska, J., Pravda, L., Vařeková, R.S. and Thornton, J.M. (2018) PDBsum: Structural summaries of PDB entries. *Protein science*, **27**, 129-134.
